# Bacterial Form I’ rubisco has smaller carbon isotope fractionation than its Form I counterpart

**DOI:** 10.1101/2023.03.01.530675

**Authors:** Renée Z. Wang, Albert K. Liu, Douglas M. Banda, Woodward W. Fischer, Patrick M. Shih

## Abstract

Form I rubiscos evolved in Cyanobacteria ≥2.5 billion years ago and are enzymatically unique due to the presence of small subunits (RbcS) that cap both ends of an octameric large subunit (RbcL) rubisco assembly to form a hexadecameric (L_8_S_8_) holoenzyme. Although RbcS was previously thought to be integral to Form I rubisco stability, the recent discovery of a closely related sister clade of octameric rubiscos (Form I’; L_8_) demonstrates that the enzyme complex assembles without small subunits (Banda et al. 2020). Rubisco also displays a kinetic isotope effect (KIE) where the 3PG product is depleted in ^13^C relative to ^12^C. In Cyanobacteria only two Form I KIE measurements exist, making interpretation of bacterial carbon isotope data difficult. To aid comparison, we measured *in vitro* the KIEs of Form I’ (*Candidatus* Promineofilum breve) and Form I (*Synechococcus elongatus* PCC 6301) rubiscos and found the KIE to be smaller in the L_8_ rubisco (16.25 ± 1.36‰ vs. 22.42 ± 2.37‰ respectively). Therefore, while small subunits may not be necessary for protein stability, they may affect the KIE. Our findings may provide insight into the function of RbcS and allow more refined interpretation of environmental carbon isotope data.

## 1. Introduction

Rubisco is a keystone enzyme linking Earth’s inorganic and organic carbon cycles, which makes it a prime target for bioengineering with regard to food systems and carbon sequestration. It is the most abundant protein on Earth today (1) because it catalyzes the essential carbon fixation step in one of the most ecologically dominant carbon-fixing metabolisms, the Calvin Benson Basshom (CBB) cycle in oxygenic photosynthesis (2)). Rubisco and oxygenic photosynthesis form the basis of our food web in terrestrial and marine systems because both eukaryotic and bacterial primary producers utilize rubisco to convert inorganic carbon (CO_2_ and HCO_3_^-^) into biomass that is then consumed by heterotrophs up the food chain. In addition, the annual flux of CO_2_ fixed by rubisco is very large, representing a significant sink in the global carbon cycle. Gross primary productivity (GPP), which accounts for all forms for carbon fixation but is vastly dominated by oxygenic photosynthesis, is ≈120 Gt C yr^-1^ in terrestrial (3) and ≈100 Gt C yr^-1^ in marine environments (1,4), compared to ≈10 Gt C yr^-1^ emitted of anthropogenic fossil CO_2_ (5). Therefore, multiple efforts exist to engineer a ‘better’ rubisco that fixes more CO_2_ in order to increase crop yields and sequester anthropogenic CO_2_, among many other motivations (see (6) for review).

However, these bioengineering approaches are driven by our current understanding of rubisco’s evolutionary history, which itself is based on our understanding of past Earth environments. These evolutionary questions largely center on the canonical paradox that, despite being a central metabolism enzyme, rubisco is: i) ‘slow,’ and ii) ‘confused’ because it can fix O_2_ instead of CO_2_ (7), which invokes a salvage pathway that costs ATP, reducing power, and carbon (8). This paradox is usually solved by considering the atmospheric composition when rubisco first evolved more than 2.5 billion years ago, when CO_2_ was much higher (potentially up to ≈20x present atmospheric levels in the Precambrian (9)) and O_2_ only existed at trace levels (2). But, in a Shakespearean tragedy, once rubisco was incorporated into the greater metabolism of oxygenic photosynthesis, it poisoned the very world it came from – successful CO_2_ fixation was coupled with oxygenation that permanently changed the atmosphere to where O_2_ is dominant (≈20%) and CO_2_ is trace (≈0.04%) today. Now saddled with a rubisco evolved from a chemical world that no longer exists, diverse land plants, algae, and Cyanobacteria have independently evolved complex CO_2_ concentrating mechanisms (CCMs) that effectively hyper-concentrate CO_2_ at the expense of O_2_ around rubisco (10) – in effect, replicating the ancient atmosphere within their own cells. To accommodate the low carboxylation rate, those without CCMs (like C3 plants) instead produce this enzyme at such high concentrations that up to 65% of all soluble protein in leaf extracts is just rubisco (11). This narrative, contingent on our understanding of the geologic carbon cycle, suggests either that rubisco is an ‘accident’ of evolutionary history, or that it is truly the optimal enzyme designed by evolution for a difficult task. If the former is true, then engineering efforts to build a better rubisco are reasonable; if not, then these efforts are a folly. Therefore, a better understanding of the evolutionary history of this enzyme is critical to rubisco engineering efforts.

Rubisco is also notable because it displays a strong carbon kinetic isotope effect (KIE) where it preferentially fixes ^12^CO_2_ over ^13^CO_2_ due to the rate of carboxylation being slightly faster for ^12^CO_2_ (12). This effect is typically reported in delta (δ^13^C) and epsilon (ε) notation in units of per mille (‰), where δ^13^C = [^13^R_sa_ / ^13^R_ref_ - 1]*1000 and ^13^R is the ratio of ^13^C/^12^C in the sample or reference respectively. ε is roughly the difference in δ^13^C between the product and the reactant (ε_Rubisco_ ≈ δ^13^C_3PG_ - δ^13^C_CO2_). Thirteen unique rubisco KIEs (ε_Rubisco_ values) have been measured across a limited range of phylogeny and species, but measurements so far suggest that rubisco fractionates at roughly 20-30‰ (for recent review see (13)).

This KIE is useful because it allows us to track mass flux through complex systems in both modern and ancient environments (14), and because it may give insight into non-isotopic enzyme kinetics (15). Since all biomass is ultimately synthesized from 3PG in autotrophs utilizing the CBB cycle, rubisco’s KIE is inherited by bulk biomass such that organic carbon is also relatively depleted in ^13^C relative to inorganic carbon. Therefore, when incorporated into larger metabolic models of carbon fixation, rubisco KIEs have facilitated the estimation of water use efficiency in plants (16), the efficiency of carbon fixation in bacterial and eukaryotic algae (17), the contribution of terrestrial plants to global GPP (18), and the proportion of C3 vs. C4 plants in mammalian diets (19) among many other examples. Similarly, in ancient environments, it has been used to estimate paleo atmospheric CO_2_ levels (20, 21), track the inorganic and organic carbon cycle through time (22), and the diet of ancient mammals (23). In addition, rubisco KIEs have been used to support interpretation of important non-isotopic kinetic parameters like the inverse correlation between specificity for CO_2_ over O_2_ (S_C/O_) and rate of carboxylation (V_C_)(15). Therefore, knowing the KIEs of many rubiscos is valuable because it facilitates empirical measurements of mass flux in many systems, natural and engineered, where other measurements may be difficult.

However, the landscape of rubisco evolution and its effect on KIE has not been well characterized. This is particularly true in Cyanobacteria, the organism within which rubisco and oxygenic photosynthesis is thought to have evolved. Most rubisco KIEs have been measured for Form IB rubiscos from plants, and in Cyanobacteria, only one Form IA and one Form IB rubisco KIE has been measured ((24, 25), for recent review see (13)). This is particularly important for reconstructing paleo pCO_2_ levels because direct measurements of the atmosphere from ice core records only extend back ≈1 million years (26), so for the remainder of Earth’s 4.6 billion year history we must rely on indirect measurements such as the carbon isotope record, globally assembled measurements of δ^13^C in the inorganic and organic carbon bearing phases of sedimentary rocks (27). Interpretation of these records relies on geochemical models, largely based on modern organisms, that incorporate the rubisco KIE to explain most of the offset in δ^13^C between inorganic and organic carbon pools (see (28) for recent review of current models). These models inform our understanding of paleo atmospheres which then influence our ideas of rubisco evolution in the past and engineering strategies in the present. It is therefore critical that we better understand the evolution of rubisco’s KIE through time because it underlies many assumptions we have about interpreting both the past and present.

We therefore tried to address this gap in knowledge by studying one key example, a Form I rubisco that lacks the small subunit. All forms of rubisco are assembled from the basic functional building block of dimers (L_2_), where two large subunits (RbcL) are assembled head-to-tail. This is the smallest known catalytically active form of rubisco. Form I rubiscos, the most ecologically abundant form of the enzyme, are hexadecameric holoenzymes (L_8_S_8_) composed of four dimers with eight small subunits (RbcS) that cap both ends of the junction between adjacent dimers. The small subunit is unique to Form I rubiscos, so it has traditionally been thought that RbcS was integral to both Form I protein stability and its evolutionary history (29). However, a novel clade of rubiscos lacking small subunits sister to Form I has recently been discovered through metagenomic analyses, and a representative octameric rubisco (L_8_) was successfully purified and kinetically characterized (30). This clade, Form I’, likely diverged before the evolution of Cyanobacteria and the small subunit (30); therefore, studying rubiscos from this clade presents a unique opportunity to study the effect of evolution on rubisco KIEs. We therefore measured *in vitro* the KIE of an L_8_S_8_ Form I rubisco from *Synechococcus elongatus* PCC 6301 in comparison to the KIE of an L_8_ Form I’ rubisco from *Candidatus* Promineofilum breve. We found the fractionation to be smaller in the L_8_ rubisco compared to the L_8_S_8_ rubisco (16.25 ± 1.36‰ vs. 22.42 ± 2.37‰ respectively). Our results imply that while the presence of a small subunit is not necessary for protein function, it may affect the KIE. Our findings may help provide insight into the function of the small subunit and allow more refined interpretation of carbon isotope data in environments, past and present, where Form I’ rubiscos may be unknowingly operating.

## 2. Materials and Methods

### 2.1 Delta Notation (δ^13^C)

Carbon isotope data were reported using delta notation (δ^13^C) in units of per mille (‰) where δ^13^C = [^13^R_sa_ / ^13^R_ref_ - 1]*1000, where the subscripts ‘sa’ and ‘ref’ denote sample and reference respectively and ^13^R is the ratio of ^13^C/^12^C. All values in this study were reported relative to the Vienna Pee Dee Belemnite (VPDB) reference.

### 2.2 Rubisco Purification

The rubiscos used here were purified according to previous methodologies and had their kinetics characterized previously (30, 31). Briefly, 14xHis-bdSUMO-tagged *Candidatus* P. breve rubisco and untagged *S. elongatus* PCC 6301 rubisco were expressed in BL21 DE3 Star *E. coli* cultures. P. breve enzyme was prepared by conducting Ni-NTA affinity purification on clarified lysate, followed by subsequent purification by anion exchange chromatography and size exclusion chromatography. *Syn*6301 enzyme was prepared by subjecting clarified lysate to ammonium sulfate precipitation at the 30-40% cut, followed by subsequent purification by anion exchange chromatography and size exclusion chromatography. The enzyme was then stored on dry ice and the KIE assay performed within one week. For Figure 1, UCSF ChimeraX was used for visualization of protein models and preparation of manuscript figures (32, 33).

**Figure 1:**
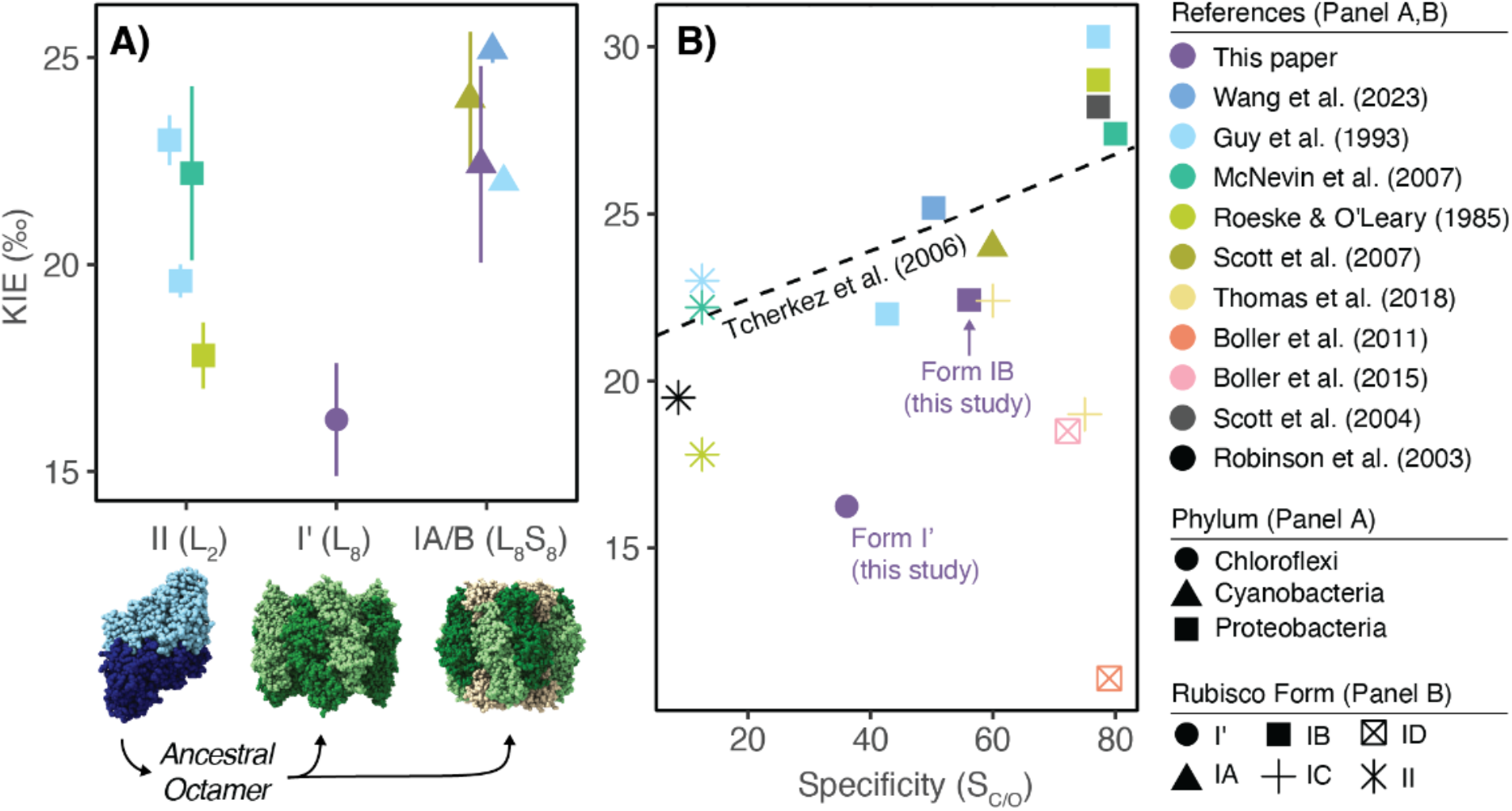
Form I’ rubisco fractionates less than both Form II and Form I rubiscos, and cannot be explained by prior model relating specificity and KIE. A) KIE (‰) for relevant bacterial Form II (L_2_), Form I’ (L_8_), and Cyanobacterial Form IA/B (L_8_S_8_) rubiscos with representative rubisco structures below; Protein Data Bank (PDB) codes from left to right: 5RUB, 6URA, 1RBL. Hypothesized evolutionary pathway is shown in black arrows, showing that ancestral dimers (L_2_) likely evolved to a common ancestral octamer (L_8_) (43) that then speciated into either Form I’ (L_8_) or Form I (L_8_S_8_) rubiscos (30). Rubisco phylum is shown as shapes and references are shown in colors. All Form II KIEs are from *Rhodospirillum rubrum* (25, 44, 45), Form I’ measurement is from *Candidatus* Promineofilum breve (this study), all Form IB rubiscos are from *Synechococcus elongatus* PCC6301 or 7942 (identical RbcL and RbcS sequence) (25, 46) and this study, Form IA KIE is from *Prochlorococcus marinus* MIT9313 (24). Error is reported as 95% confidence intervals for (24); as standard deviation for this study and (44–46); as standard error for (25).See Supplemental Table S2 for literature values used, notes on variation between measurements, and rationale for which data was included and excluded. For recent compilation of all measured rubisco KIEs, see (13). B) Compilation of additional KIE and specificity values in Form IC and ID rubiscos, in addition to data shown in Figure 1A. Forms shown in shapes, references shown in same colors as in Panel A. Panel B is based on Figure 4 in (36). See Table S2 and S3 for compilation of data used. Dotted line indicates original linear regression from Figure 3F in (15).

### 2.3 Rubisco KIE Assay Summary

We used a substrate depletion method to measure the KIE of each rubisco as used previously in similar studies (25, 34–36). Briefly, this method relies on measuring the time-varying δ^13^C value of the CO_2_ pool as the reaction goes to completion instead of directly measuring the difference in δ^13^C between the initial CO_2_ and final 3PG pool. The KIE is then calculated from these data using a Rayleigh relationship, which considers the log-log transformation of the CO_2_ isotope data against the fraction of substrate consumed.Linear regression of this data can then be converted to a measure of the instantaneous isotope fractionation—the empirical measure of the isotope effect associated with rubisco carboxylation. With this formulation larger KIEs correspond to steeper slopes in a Rayleigh Plot.

The assay mix we used is based on previous similar studies. In this set-up, inorganic carbon is supplied as HCO_3_^-^ which is converted to CO_2_ by a carbonic anhydrase (CA), typically derived from bovines. CO_2_ and RuBP is then catalyzed by rubisco to create 3PG. Therefore, our reaction mixture contains CA, rubisco, HCO_3_^-^ and RuBP to yield the full reaction, and additional reagents like: i) MgCl_2_ to support correct rubisco active site metallation, ii) bicine as a buffer, iii) dithiothreitol (DTT) to prevent rubisco oxidation and degradation (37).

In our experiment, instead of limiting CO_2_, we limited RuBP. In addition, *f* (the proportion of CO_2_ remaining) is typically known from an external measurement. Prior experiments have labored to constrain *f* by taking a separate aliquot of the assay to measure CO_2_ concentration directly (25, 35). In our experiment, we converted sampling time to *f* by fitting our data to the model y = a*EXP(-b*x)+c based on the fact that the δ^13^C of the reactant pool will increase during the reaction and then asymptote to a fixed value as the reaction ceases (i.e. no further carbon isotope discrimination can occur because rubisco can no longer pull from the CO_2_ pool as RuBP runs out). In essence, we are interested in the curvature of this line, similar to prior rubisco assays where the δ^13^C of the reaction vessel headspace was monitored continually on a membrane inlet mass spectrometer (34) instead of traditional methods where discrete aliquots are taken (25). See below and Supplemental for further discussion.

### 2.4 Assay Preparation and Execution

Prior to running the KIE assay, the activity of bovine erythrocytes CA (Sigma Aldrich; C3934) was checked following manufacturer guidelines (38) We found a value of 3,368 W-A units / mg protein, which exceeded the product stated value of ≥2,OOO W-A units / mg protein and so proceeded to use this active CA enzyme prep in the KIE assay.

Glass sampling vials with septum tops (‘Exetainer,’ 12 mL, Labco) were prepared. First, three external standards were prepared by weighing out Carrara marble standards (CIT_CM2013, δ^13^C = 2.0 ± 0.1‰) into individual exetainers. Standards were then sealed within each tube, purged with He gas for 5 minutes, and then acidified by needle injection with concentrated phosphoric acid (42% v/v). Then, three HCO_3_^-^ substrate exetainers were also sealed, purged with He gas, acidified by needle injection of phosphoric acid to convert HCO_3_^-^ to CO_2_, and placed in a 70°C water bath for at least 20 minutes. Finally, 22 exetainer sampling vials were prepared for the rubiscos (12 for L_8_, 10 for L_8_S_8_). All sampling tubes were first sealed and purged with He gas for 5 minutes, and then injected with ~1 mL of anhydrous phosphoric acid. The phosphoric acid both stops the reaction progress, and converts all dissolved inorganic carbon species into CO_2_ for analysis.

Next, the reaction assay for each rubisco was prepared. First, a CA stock solution was made by dissolving bovine erythrocytes CA into DI water. Next, an RuBP stock solution was made by dissolving D-Ribulose 1,5-bisphosphate sodium salt hydrate (Sigma Aldrich; R0878) in DI water. Then, one drop of concentrated hydrochloric acid (38% v/v) was added to 20 mL of autoclaved DI water while it was simultaneously stirred with a stir bar and vigorously bubbled with N_2_ gas for 10 minutes to remove any residual HCO_3_^-^ or CO_2_. Then, while N_2_ gas was blown over the surface of the solution to inhibit O_2_, reagents were added to create a final concentration of 100 mM bicine, 30 mM MgCl_2_, 1 mM dithiothreitol (DTT), and 6.25 mM NaHCO_3_. pH was adjusted to 8.5 with NaOH and HCl. CA from the CA stock was added, and then either the L_8_ or L_8_S_8_ rubisco was added to the solution. 0.996 mg of L_8_S_8_ and 1.18 mg of L_8_ rubisco were used. The solution was gently bubbled with N_2_ gas for 10 minutes while rubisco ‘activated.’ While the solution was bubbling, the syringes used for each rubisco assay were rinsed with ethanol and water. We used a separate 25 mL gas-tight syringe with a sample-locking needle for each rubisco (Ref #86326, Model 1025 SL SYR, Hamilton Company).

RuBP was then added to each reaction assay and mixed through pipetting and swirling. This entire solution was then quickly transferred to a gas-tight syringes. The first time point (t=0 min) was taken as quickly as possible after transfer. To sample, ~1 mL of the reaction assay was injected into the pre-prepared sampling exetainer containing phosphoric acid. Each assay was sampled 10-12 times over 390 minutes.

A control was run in a separate experiment, where all the assay components were mixed together with the exception of a rubisco enzyme. The δ^13^C of the measured headspace did not change appreciably during this time period, with δ^13^C = −0.42 ± 0.03‰ at 0 minutes and δ^13^C = −0.55 ± 0.03‰ at 277 minutes. The absolute values of these measurements reflect the δ^13^C of the substrate used on that experimental day and cannot be related to the data shown here.

### 2.5 Isotopic Measurement

The δ^13^C of CO_2_ in the headspace of each exetainer was measured on a Delta-V Advantage with Gas Bench and Costech elemental analyzer at Caltech. Before measuring samples, two tests were performed to ensure the instrument was functioning normally: i) An ‘on/off’ test with an internal CO_2_ standard for instrument sensitivity and to establish a baseline intensity at a ‘zero’ CO_2_ concentration, and ii) A linearity test where the concentration of CO_2_ was increased linearly within the designated sensitivity range of the instrument to ensure that a linear increase in CO_2_ concentration corresponds to a linear increase in electrical signal on the collector cups. We measured the concentration of ^12^CO_2_ at mass 44, and ^13^CO_2_ at mass 45. The instrument was also tuned to ensure that each mass was measured at the center of its mass peak.

The headspace of each sample and standard was measured 10 times (10 analytical replicates), with an internal CO_2_ reference run before and after each suite of 10 analytical replicates. Data was visually inspected to ensure the sample was being measured within the correct sensitivity range of the instrument (i.e. of similar intensity and pressure as the internal CO_2_ reference). The ‘raw’δ^13^C values were then corrected relative to VPDB using the three external standards. Assay results can be seen in Table S1 and Figure S1A.

### 2.6 Calculation of KIE

We first pre-processed the data by assessing which data points to fit. We expect the δ^13^C of CO_2_ to increase following an exponential curve that eventually reaches an asymptote, but the last few data points start to decrease in δ^13^C. This may be due to a variety of reasons, including: 1) Ambient CO_2_ contaminating the exetainer containers as they are left out after the reaction; 2) Re-equilibration of the aqueous and gaseous inorganic carbon pools; 3) Instrument error upon needle sampling of exetainer vial. Because exponential curves are linear in a log-log space, we therefore log-transformed the data points then systematically fit a linear regression through varying sets of data and calculated the resulting error (adjusted R^2^ value). The adjusted R^2^ value consistently decreased after data point 9 for L_8_, and after data point 8 for L_8_S_8_ (Figure S1B,C). Therefore, we proceeded to use data points 1-9 for L_8_ and 1-8 for L_8_S_8_.

We then converted time to *f*, the fraction of the inorganic C pool remaining. Since RuBP was the limiting substrate, we could calculate the moles of CO_2_ consumed if we assume: i) A 1:1 ratio of RuBP to CO_2_ was utilized by Rubisco, and ii) Full consumption of the RuBP pool. For each rubisco assay, 125 μmol of RuBP and 9.84 μmol of NaHCO_3_ were added. Therefore, 7.87% of the initial CO_2_ pool was consumed, or *F* = 0.9213. We then assume that *f* = 1 at *t* = 0, and *f* = 0.9213 at the upper bound of the fit. A general model of y = a*EXP(-b*x)+c was applied to the data, though with carbon isotope data in the ^13^R format instead of the δ^13^C format because ^13^R values can be manipulated arithmetically while δ^13^C values cannot (39). The model was then fitted using the non-linear least squares function (call: *n/s*(); R Statistical Software (v4.1.0; R Core Team 2021, (40))). Model outputs are shown in Table S4 and Figure S2.

Time was then converted to *f* using the equation:

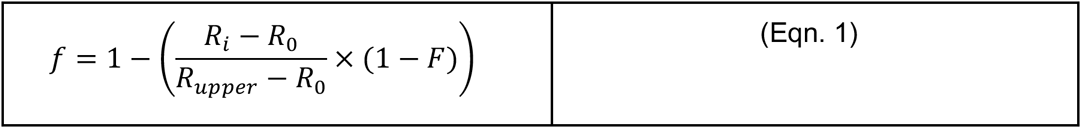

Where *R_0_* is the first measured ^13^R value in each set of data, *R_upper_* is the fitted parameter *c* from and *F* = 0.9213, which is calculated from the amount of RuBP added to the assay.

Next, a correction was done to account for the C isotope fractionation between CO_2_ and HCO_3_^-^ at equilibrium, where CO_2_ is ~8‰ lighter (more negative δ^13^C value) than HCO_3_^-^ (41). We followed the correction outlined in (25) where the adjustment is applied before linear regression in a Rayleigh plot:

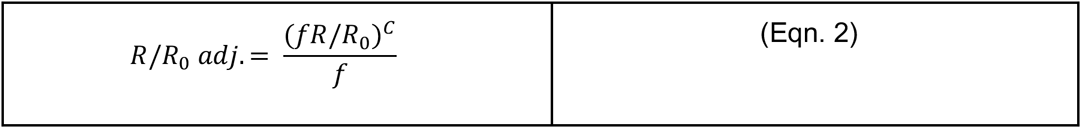

Where C = (1.009 + 10^(pK-pH))/(1+10^(pK-pH)). The pK is that of carbonic acid, for which we used a value of 6.4 (42). The pH of the L_8_S_8_ assay was 8.49, and the pH of the L_8_ assay was 8.52.

Finally, a Rayleigh Plot was made for each rubisco plotting ln(^13^R/^13^R_0_)_adj._*1000 vs. -ln(*f*) (Figure S3). The best fit slope, *D*, was calculated using a linear regression (call: *lm();* R Statistical Software (v4.1.0; R Core Team 2021, (40))). *D* was then converted to Δ, the KIE, using the equation Δ =*D*/(1-*D*/1000) (25). Doing so, we found the KIE of the L_8_S_8_ rubisco to be 22.42 ± 2.37, and 16.25 ± 1.36 for the L_8_ rubisco. Results are shown in Table 1.

**Table 1:** KIE and non-isotopic kinetic measurements from L_8_ vs. L_8_S_8_ rubiscos. KIEs were measured in this study using the substrate depletion method (25, 34–36); see Methods for more detail. Non-isotopic kinetic measurements are from (30). V_C_ and V_O_ indicate maximum carboxylation and oxygenation rates under substrate-saturated conditions, respectively; K_C_ and K_O_ are Michaelis constants for the carboxylation and oxygenation reactions, respectively; S_C/O_ indicates specificity, a unitless measure of the relative preference for CO_2_ over O_2_ and is calculated as S_C/O_ = (V_C_/K_C_)/(V_O_/K_O_). Uncertainties on non-isotopic kinetics reflect mean ± s.e.m. from multiple experiments; see (30) for more detail. Error on KIEs reflect mean ± s.d. from model fitting uncertainty from one experiment; see Methods and Supplemental for more detail.

| Strain | Rubisco | KIE (‰) | $V_c$ ($s^{-1}$) | $K_c$ ($\mu\text{M}$) | $S_{c/o}$ | $V_o$ ($s^{-1}$) | $K_o$ ($\mu\text{M}$) |
| --- | --- | --- | --- | --- | --- | --- | --- |
| *Synechococcus elongatus* PCC6301 | $L_8S_8$ | $22.42 \pm 2.37$ | $14.3 \pm 0.71$ | $235 \pm 20.0$ | $56.1 \pm 1.3$ | 1.10 | $983 \pm 81$ |
| *Candidatus*<br>Promineofilum breve | L<sub>8</sub> | 16.25 ±<br>1.36 | 2.23 ±<br>0.04 | 22.2 ± 9.7 | 36.1 ± 0.9 | 1.11 | 401 ± 115 |

## 3. Results

### 3.1 L_8_ rubisco has smaller KIE than its L_8_S_8_ counterpart

The KIE of the L_8_ rubisco is ≈5‰ less than that of the L_8_S_8_ rubisco (16.25 ± 1.36‰ vs. 22.42 ± 2.37‰ respectively). We note that there is variation among KIE measurements of similar or the same strains. Prior measurements which we compare our data against (Figure 1A, Table S2) are bacterial (Form II, Form I’) or Cyanobacterial (Form I) rubisco measurements where a pure enzyme, substrate-depletion assay like ours was done from well-characterized strains where rubisco was obtained through expression and subsequent purification from *E. coli.*Therefore, we did not include measurements where a non-native bacterial rubisco was expressed by another organism *in vivo* and KIE calculated by extrapolating ratios of intracellular to extracellular CO_2_ (47), measurements where the strain cannot be cultured separately from a host organism (48), nor measurements from plants or the *Solemya velum* symbiont because it is not a member of the Cyanobacteria (35). It has been proposed and measured that rubisco KIEs vary with pH, temperature, metal ion concentrations (49, 50), yet other studies contradict this claim (51) and have instead proposed that much of the variation in the literature reflects experimental uncertainties rather than intrinsic variations in KIE (16). This study and (46) measured an L_8_S_8_ rubisco KIE from *Synechococcus elongatus* PCC6301 and 7942 respectively (identical RbcL and RbcS sequences) in similar assay conditions but found values that are similar but do not overlap in uncertainty, supporting the conclusion that variations in reported KIE values is due to experimental uncertainty rather than intrinsic enzymatic variations, However, the KIEs presented in Figure 1 were measured in assays that span a range of pH, temperature, and MgCl_2_ concentration (Table S2), notably with increasing MgCl_2_ concentration corresponding with increasing KIEs measured in the Form II rubisco by (25). Because of the lack of repeated, rigorous measurements of multiple rubisco KIEs across variations relevant parameters (i.e. pH, temperature, metallation), it is difficult to conclude what is causing the variation in KIE values across studies. Therefore, we can only conclude that the L_8_ rubisco KIE is less (by roughly 5‰) than its L_8_S_8_ counterpart measured in this study, and the range of L_8_S_8_ rubiscos measured from previous studies.

Similarly, compared to prior Form II (L_2_) rubisco KIE measurements, the Form I’ (L_8_)rubisco may fractionate less. Compared to Form I KIEs, there is wider variation in previously measured Form II KIEs, with the Form I’ rubisco measured here overlapping in value with one Form II rubisco within uncertainty (45). We note that all the Form II data presented here are from one species, *Rhodospirillum rubrum*, though the specific strain is not reported for all studies. Therefore, the variations may reflect experimental uncertainty with the exception of the (25) measurement where MgCl_2_ concentration was changed. Therefore, we are not confident concluding either way if the L_8_ KIE is less than the L_2_ KIE or not.

## 4. Discussion

### 4.1 Presence or absence of RbcS external to active site may influence KIE

Rubisco KIEs have also been used to support conclusions gleaned from non-isotopic kinetic parameters, both to better understand the reaction mechanism and to offer complementary data to traditional measurements, but our results belie an easy interpretation within that existing framework. The dominant theory in this field posits that rubisco specificity is positively correlated with the CO_2_ KIE because of an observed increase in carbon isotope fractionation, but not oxygen isotope fractionation, with specificity (15, 25). This argument originates from studies of deuterium (D) isotope effects on enzymatic reaction rates, which have been traditionally performed because deuterium displays a much larger (and easier to measure) KIE due to the large relative mass difference between D and its major isotope, H, in comparison to other rare isotopes like ^13^C vs. ^12^C or ^15^N vs. ^14^N (52). These foundational experiments have led to the conclusion that the isotope effect is determined at the rate-limiting step at the transition state, and small asymmetries in the transition state caused by transition state structure will cause small variations in the isotope effect (52, 53). Applied to rubisco, (15) proposed that the inherent difficulty in binding a ‘featureless’ CO_2_ vs. O_2_ molecule has caused natural selection in the transition state, where rubiscos that maximize the structural difference in transition states for carboxylation vs. oxygenation are able to be more specific. That then causes a trade-off where greater resemblance to the final carboxyketone product causes the product to also be tightly bound, leading to a higher S_C/O_ correlating with a lower V_C_, but also a prediction that the intrinsic KIE for CO_2_ addition (but not O_2_ addition) should increase as the transition state becomes more product like - i.e., higher specificity rubiscos should have higher KIEs - which is indeed what the data at the time support (15). This has also led to the conclusion that rubisco is actually perfectly optimized for the time and places where it is found today, precluding any opportunity to use rubisco engineering to achieve increased biomass yields(15).

However, new CO_2_ KIE measurements that do not show a correlation with specificity are empirically questioning this conclusion (Figure 1B). Prior studies (36) have pointed out that the spread in KIE data, particularly at high specificity, cannot easily be described by a simple inverse relationship or linear regression. Indeed, our Form I’ measurement lies below the original regression line proposed in (15) – its KIE is effectively too low given what one would predict via its specificity. However, although an increasing spread in CO_2_ KIE becomes apparent as more rubiscos are measured, they cannot directly address the dominant theory because of the general dearth of O_2_ KIE measurements. In addition, specificity is typically not reported in the same study with KIE (see notes in Table S2, S3), so some of the spread in Figure 1B may be due to uncertainties in the true specificity for the given rubisco measured. Therefore, additional paired measurements of CO_2_ and O_2_ KIEs with specificity are necessary before a new theory relating isotopic and non-isotopic kinetics can be proposed; more data is needed to decide between potential theories.

In addition, this transition state optimization theory is based on the assumption that it is the active site (which binds the intermediary carboxylation or oxygenation product) that concurrently affects both specificity and KIE, so the naive assumption is that the absence or presence of the small subunit, which does *not* contain the active site, should not affect KIE. Unexpectedly, the L_8_ rubisco fractionates roughly 5‰ less than that of the L_8_S_8_ rubisco (16.25 ± 1.36‰ vs. 22.42 ± 2.37‰ respectively). The specificity of the L_8_ rubisco is indeed less than that of the L_8_S_8_ (36.1 ± 0.9 vs. 56.1 ± 1.3 respectively, (30)) but this may be a coincidence because that prediction is based on a theory reliant on rubisco’s active site which the small subunit does not directly impact. Our comparative study suggests the tantalizing hypothesis that the small subunit increases rubisco KIEs. One way to test this hypothesis is to strip RbcS from a Form I rubisco, but this is not immediately doable because doing so makes the Form I rubisco inactive, so far. Fortunately, learning more about Form I’ rubisco may make this future experiment possible. Alternately, it has been shown that distal mutations from the active site affecting oligomerization can affect enzyme kinetics, which is somewhat analogous to losing RbcS that does not directly interact with the active site – KIE measurements from such rubiscos may shed help shed light on the relationship between RbcS, specificity, and KIE (54). In addition, dual CO_2_ and O_2_ KIE measurements of other novel Form I’ rubiscos compared to Form I rubiscos, across a range of assay parameters, would be needed for a more robust comparative study. Therefore, it remains an open question what structural and biochemical aspects of rubisco may also affect KIEs in addition to active site and transition state theory mechanisms.

### 4.2 Supports prior work positing that rubisco KIEs vary across phylogeny in the modern

Our work supports previous work showing that the rubisco KIE varies across phylogeny in the modern, though with the caveat that few unique rubiscos have been measured, there is variation across experiments, and the vast majority of measurements are from Form I rubiscos (Figure 1B, and see (36, 55) for recent compilation across phylogeny). A smaller KIE measured from one novel Form I’ rubisco, in comparison to the bacterial Form I rubiscos, supports this observation, though more measurements across the Form I’ clade are needed to quantify any potential in-clade variation.

### 4.3 Supports prior work positing that rubisco KIEs may have varied across time, in addition to phylogeny

If we view L_8_ as an evolutionary ‘missing link’ between the evolution of L_2_ and L_8_S_8_ rubiscos, this measurement supports the idea that rubisco KIE may have varied across evolutionary time. Prior work has explored this question by measuring the KIE of a putative Precambrian ancestral Form IB rubisco reconstructed using a combination of phylogenetic and molecular biology techniques (56); that work found the ancestral rubisco to fractionate less than its modern counterpart (17.23 ± 0.61‰ vs. 25.18 ± 0.31‰ respectively) (46). Interestingly, the Form I’ and putative ancestral Form IB rubisco have similar, lower KIE values (16.25 ± 1.36‰ vs. 17.23 ± 0.61‰ respectively) compared to most modern Form I rubiscos (roughly 20-30‰ (for recent review see ((13)). This supports prior predictions that the KIE should have varied over geologic time in response to changing pCO_2_, though that prediction was based on an assumption of inverse correlation between specificity (selected for by changing CO_2_/O_2_ levels) and KIE (15). This implies that the KIE of ancestral rubiscos may have been lower than modern rubiscos today, though this is a tentative hypothesis that, by necessity, relies on ancestral enzyme reconstruction and comparative biology techniques instead of direct measurements of ‘true’ ancestral enzymes.

Finally, it is hypothesized that the small subunit may have evolved in response to rising atmospheric oxygen levels roughly 2.4 billion years ago because the high V_O_ stabilization that RbcS offers allows simultaneous exploration of RbcS and RbcL protein space (30). Therefore, understanding the KIE of Form I’ rubiscos may allow us to better understand changes in rubisco biochemistry that may have accompanied evolutionary changes and facilitate better tracking of carbon mass flux at key times in Earth’s evolutionary history.

## Data Availability Statement

All data used in this study are presented in the supplement.

## References

1. Y. M. Bar-On, R. Milo, The global mass and average rate of rubisco. Proc Natl Acad Sci USA 116, 4738–4743 (2019).

2. W. W. Fischer, J. Hemp, J. E. Johnson, Evolution of oxygenic photosynthesis. Annu. Rev. Earth Planet. Sci. 44, 647–683 (2016).

3. C. Beer, et al., Terrestrial gross carbon dioxide uptake: global distribution and covariation with climate. Science 329, 834–838 (2010).

4. C. B. Field, M. J. Behrenfeld, J. T. Randerson, P. Falkowski, Primary production of the biosphere: integrating terrestrial and oceanic components. Science 281, 237–240 (1998).

5. P. Friedlingstein, et al., Global carbon budget 2022. Earth Syst. Sci. Data 14, 4811–4900 (2022).

6. R. J. Spreitzer, M. E. Salvucci, Rubisco: structure, regulatory interactions, and possibilities for a better enzyme. Annu. Rev. Plant Biol. 53, 449–475 (2002).

7. G. H. Lorimer, T. J. Andrews, Plant photorespiration—an inevitable consequence of the existence of atmospheric oxygen. Nature 243, 359–360 (1973).

8. T. J. Andrews, G. H. Lorimer, The Biochemistry of Plants: A Comprehensive Treatise, Vol. 10, Photosynthesis, M. D. Hatch, N. K. Boardman, Eds. (1987).

9. N. D. Sheldon, Precambrian paleosols and atmospheric CO2 levels. Precambrian Res. 147, 148–155 (2006).

10. A. Flamholz, P. M. Shih, Cell biology of photosynthesis over geologic time. Curr. Biol. 30, R490–R494 (2020).

11. R. J. Ellis, The most abundant protein in the world. Trends Biochem. Sci. 4, 241–244 (1979).

12. G. D. Farquhar, J. R. Ehleringer, K. T. Hubick, Carbon Isotope Discrimination and Photosynthesis. Annu. Rev. Plant Physiol. Plant Mol. Biol. 40, 503–537 (1989).

13. A. K. Garcia, C. M. Cavanaugh, B. Kacar, The curious consistency of carbon biosignatures over billions of years of Earth-life coevolution. ISME J. 15, 2183–2194 (2021).

14. J. M. Hayes, Fractionation of carbon and hydrogen isotopes in biosynthetic processes. Reviews in Mineralogy and Geochemistry 43, 225–277 (2001).

15. G. G. B. Tcherkez, G. D. Farquhar, T. J. Andrews, Despite slow catalysis and confused substrate specificity, all ribulose bisphosphate carboxylases may be nearly perfectly optimized. Proc Natl Acad Sci USA 103, 7246–7251 (2006).

16. G. D. Farquhar, M. H. O’Leary, J. A. Berry, On the relationship between carbon isotope discrimination and the intercellular carbon dioxide concentration in leaves. Aust. J. Plant Physiol. 9, 121 (1982).

17. T. D. Sharkey, J. A. Berry, “Carbon Isotope Fractionation of Algae as Influenced by an Inducible CO2 Concentrating Mechanism” in Inorganic Carbon Uptake by Aquatic Photosynthetic Organisms, W. J. Lucas, J. A. Berry, Eds. (The American Society of Plant Physiologists, 1985), pp. 389–401.

18. J. Lloyd, G. D. Farquhar, 13C discrimination during CO2 assimilation by the terrestrial biosphere. Oecologia 99, 201–215 (1994).

19. T. E. Cerling, J. M. Harris, Carbon isotope fractionation between diet and bioapatite in ungulate mammals and implications for ecological and paleoecological studies. Oecologia 120, 347–363 (1999).

20. C. R. Witkowski, J. W. H. Weijers, B. Blais, S. Schouten, J. S. Sinninghe Damsté, Molecular fossils from phytoplankton reveal secular PCO2 trend over the Phanerozoic. Sci. Adv. 4, eaat4556 (2018).

21. R. R. Bidigare, et al., Consistent fractionation of^13^ C in nature and in the laboratory: Growth-rate effects in some haptophyte algae. Global Biogeochem. Cycles 11, 279–292 (1997).

22. M. Schidlowski, A 3,800-million-year isotopic record of life from carbon in sedimentary rocks. Nature 333, 313–318 (1988).

23. T. E. Cerling, J. M. Harris, M. G. Leakey, Browsing and grazing in elephants: the isotope record of modern and fossil proboscideans. Oecologia 120, 364–374 (1999).

24. K. M. Scott, et al., Kinetic isotope effect and biochemical characterization of form IA RubisCO from the marine cyanobacterium Prochlorococcus marinus MIT9313. Limnol. Oceanogr. 52, 2199–2204 (2007).

25. R. D. Guy, M. L. Fogel, J. A. Berry, Photosynthetic fractionation of the stable isotopes of oxygen and carbon. Plant Physiol. 101, 37–47 (1993).

26. J. A. Higgins, et al., Atmospheric composition 1 million years ago from blue ice in the Allan Hills, Antarctica. Proc Natl Acad Sci USA 112, 6887–6891 (2015).

27. J. Krissansen-Totton, R. Buick, D. C. Catling, A statistical analysis of the carbon isotope record from the Archean to Phanerozoic and implications for the rise of oxygen. Am. J. Sci. 315, 275–316 (2015).

28. E. B. Wilkes, A. Pearson, A general model for carbon isotopes in red-lineage phytoplankton: Interplay between unidirectional processes and fractionation by RubisCO. Geochim. Cosmochim. Acta (2019) https://doi.org/10.1016/j.gca.2019.08.043.

29. R. J. Spreitzer, Role of the small subunit in ribulose-1,5-bisphosphate carboxylase/oxygenase. Arch. Biochem. Biophys. 414, 141–149 (2003).

30. D. M. Banda, et al., Novel bacterial clade reveals origin of form I Rubisco. Nat. Plants 6, 1158–1166 (2020).

31. S. Saschenbrecker, et al., Structure and function of RbcX, an assembly chaperone for hexadecameric Rubisco. Cell 129, 1189–1200 (2007).

32. E. F. Pettersen, et al., UCSF ChimeraX: structure visualization for researchers, educators, and developers. Protein Sci. 30, 70–82 (2021).

33. T. D. Goddard, et al., UCSF ChimeraX: Meeting modern challenges in visualization and analysis. Protein Sci. 27, 14–25 (2018).

34. D. B. McNevin, M. R. Badger, H. J. Kane, G. D. Farquhar, Measurement of (carbon) kinetic isotope effect by Rayleigh fractionation using membrane inlet mass spectrometry for CO2-consuming reactions. Funct. Plant Biol. 33, 1115 (2006).

35. K. M. Scott, J. Schwedock, D. P. Schrag, C. M. Cavanaugh, Influence of form IA RubisCO and environmental dissolved inorganic carbon on the delta13C of the clam-chemoautotroph symbiosis Solemya velum. Environ. Microbiol. 6, 1210–1219 (2004).

36. P. J. Thomas, et al., Isotope discrimination by form IC RubisCO from Ralstonia eutropha and Rhodobacter sphaeroides, metabolically versatile members of “Proteobacteria” from aquatic and soil habitats. Environ. Microbiol. (2018) https://doi.org/10.1111/1462-2920.14423.

37. Y. Marcus, H. Altman-Gueta, A. Finkler, M. Gurevitz, Dual role of cysteine 172 in redox regulation of ribulose 1,5-bisphosphate carboxylase/oxygenase activity and degradation. J. Bacteriol. 185, 1509–1517 (2003).

38. , Enzymatic Assay of Carbonic Anhydrase for Wilbur-Anderson Units (EC 4.2.1.1) (June 8, 2022).

39. J. M. Hayes, Practice and principles of isotopic measurements in organic geochemistry. Organic geochemistry of contemporaneous and ancient sediments 5 (1983).

40. R. C. Team, R: A language and environment for statistical computing. Published online 2020. (2021).

41. W. G. Mook, J. C. Bommerson, W. H. Staverman, Carbon isotope fractionation between dissolved bicarbonate and gaseous carbon dioxide. Earth and Planetary Science Letters 22, 169–176 (1974).

42. R. E. Zeebe, D. Wolf-Gladrow, CO2 in seawater: Equilibrium, kinetics, isotopes (Elsevier, 2001).

43. L. Schulz, et al., Evolution of increased complexity and specificity at the dawn of form I Rubiscos. Science 378, 155–160 (2022).

44. D. B. McNevin, et al., Differences in carbon isotope discrimination of three variants of D-ribulose-1,5-bisphosphate carboxylase/oxygenase reflect differences in their catalytic mechanisms. J. Biol. Chem. 282, 36068–36076 (2007).

45. C. A. Roeske, M. H. O’Leary, Carbon isotope effect on carboxylation of ribulose bisphosphate catalyzed by ribulosebisphosphate carboxylase from Rhodospirillum rubrum. Biochemistry 24, 1603–1607 (1985).

46. R. Z. Wang, et al., Carbon isotope fractionation by an ancestral rubisco suggests biological proxies for CO_2_ through geologic time should be re-evaluated (2023).

47. S. von Caemmerer, Y. Tazoe, J. R. Evans, S. M. Whitney, Exploiting transplastomically modified Rubisco to rapidly measure natural diversity in its carbon isotope discrimination using tuneable diode laser spectroscopy. J. Exp. Bot. 65, 3759–3767 (2014).

48. J. J. Robinson, et al., Kinetic isotope effect and characterization of form II RubisCO from the chemoautotrophic endosymbionts of the hydrothermal vent *tubewormRiftia pachyptila. Limnol*. Oceanogr. 48, 48–54 (2003).

49. T. Whelan, W. M. Sackett, C. R. Benedict, Enzymatic Fractionation of Carbon Isotopes by Phosphoenolpyruvate Carboxylase from C4 Plants. Plant Physiol. 51, 1051–1054 (1973).

50. M. H. O’Leary, Heavy atom isotope effects in enzyme-catalyzed reactions. In’Transition States of Biochemical Processes’.(Eds R. Gandour and RL Schowen.) pp. 285–316. 285 (1978).

51. J. T. Christeller, W. A. Laing, Isotope Discrimination by Ribulose 1,5-Diphosphate Carboxylase: No Effect of Temperature or HCO(3) Concentration. Plant Physiol. 57, 580–582 (1976).

52. P. A. Frey, A. D. Hegeman, Enzymatic Reaction Mechanisms (Oxford University Press, 2007) https://doi.org/10.1093/oso/9780195122589.001.0001.

53. F. H. Westheimer, The magnitude of the primary kinetic isotope effect for compounds of hydrogen and deuterium. Chem. Rev. 61, 265–273 (1961).

54. A. K. Liu, et al., Structural plasticity enables evolution and innovation of RuBisCO assemblies. Sci. Adv. 8 (2022).

55. A. K. Garcia, et al., System-level effects of CO2 and RuBisCO concentration on carbon isotope fractionation. BioRxiv (2021) https://doi.org/10.1101/2021.04.20.440233.

56. P. M. Shih, et al., Biochemical characterization of predicted Precambrian RuBisCO. Nat. Commun. 7, 10382 (2016).

57. A. L. De Oliveira, A. Srivastava, S. Espada-Hinojosa, M. Bright, The complete and closed genome of the facultative generalist Candidatus Endoriftia persephone from deep-sea hydrothermal vents. Mol. Ecol. Resour. 22, 3106–3123 (2022).

58. D. Davidi, et al., Highly active rubiscos discovered by systematic interrogation of natural sequence diversity. EMBO J. 39, e104081 (2020).

59. K. M. Horken, F. R. Tabita, Closely related form I ribulose bisphosphate carboxylase/oxygenase molecules that possess different CO2/O2 substrate specificities. Arch. Biochem. Biophys. 361, 183–194 (1999).

60. M. R. Badger, et al., The diversity and coevolution of Rubisco, plastids, pyrenoids, and chloroplast-based CO_2_-concentrating mechanisms in algae. Can. J. Bot. 76, 1052–1071 (1998).

61. R. P. Haslam, et al., “Specificity of diatom Rubisco” in Plant Responses to Air Pollution and Global Change, K. Omasa, I. Nouchi, L. J. De Kok, Eds. (Springer Japan, 2005), pp. 157–164.

62. B. A. Read, F. R. Tabita, High substrate specificity factor ribulose bisphosphate carboxylase/oxygenase from eukaryotic marine algae and properties of recombinant cyanobacterial RubiSCO containing “algal” residue modifications. Arch. Biochem. Biophys. 312, 210–218 (1994).

63. H. J. Kane, et al., An Improved Method for Measuring the CO2/O2 Specificity of Ribulosebisphosphate Carboxylase-Oxygenase. Functional Plant Biol. 21, 449 (1994).

64. A. J. Boller, P. J. Thomas, C. M. Cavanaugh, K. M. Scott, Isotopic discrimination and kinetic parameters of RubisCO from the marine bloom-forming diatom, Skeletonema costatum. Geobiology 13, 33–43 (2015).

65. A. J. Boller, P. J. Thomas, C. M. Cavanaugh, K. M. Scott, Low stable carbon isotope fractionation by coccolithophore RubisCO. Geochim. Cosmochim. Acta 75, 7200–7207 (2011).

66. C. A. Roeske, M. H. O’Leary, Carbon isotope effects on enzyme-catalyzed carboxylation of ribulose bisphosphate. Biochemistry 23, 6275–6284 (1984).

67. H. Wickham, W. Chang, M. H. Wickham, Package ‘ggplot2’. Create elegant data visualisations using the grammar of graphics. Version 2, 1–189 (2016).

