## Supplemental Figures and Tables for "Bacterial Form I’ rubisco has smaller carbon isotope fractionation than its Form I counterpart"

### Supplementary Materials

| Rubisco | Time (min) | $\delta^{13}C$ (avg.) | $\delta^{13}C$ (std. err.) | $^{13}R$ (avg.) | $^{13}R$ (std. err.) |
|---|---|---|---|---|---|
| $L_8$ | 0 | -0.592 | 0.010 | 0.0111549 | 1.81E-07 |
| $L_8$ | 15 | -0.425 | 0.007 | 0.0111580 | 1.24E-07 |
| $L_8$ | 30 | -0.129 | 0.021 | 0.0111634 | 3.92E-07 |
| $L_8$ | 45 | 0.111 | 0.026 | 0.0111679 | 4.80E-07 |
| $L_8$ | 60 | 0.268 | 0.017 | 0.0111708 | 3.11E-07 |
| $L_8$ | 90 | 0.327 | 0.010 | 0.0111718 | 1.81E-07 |
| $L_8$ | 120 | 0.506 | 0.007 | 0.0111751 | 1.35E-07 |
| $L_8$ | 150 | 0.473 | 0.012 | 0.0111745 | 1.35E-07 |
| $L_8$ | 210 | 0.652 | 0.013 | 0.0111778 | 2.34E-07 |
| $L_8$ | 270 | 0.454 | 0.007 | 0.0111742 | 1.26E-07 |
| $L_8$ | 330 | 0.399 | 0.006 | 0.0111732 | 1.08E-07 |
| $L_8$ | 390 | 0.348 | 0.014 | 0.0111722 | 2.55E-07 |
| $L_8S_8$ | 0 | 0.599 | 0.026 | 0.0111768 | 4.68E-07 |
| $L_8S_8$ | 15 | 0.990 | 0.017 | 0.0111840 | 3.14E-07 |
| $L_8S_8$ | 30 | 1.058 | 0.008 | 0.0111853 | 1.53E-07 |
| $L_8S_8$ | 45 | 1.553 | 0.015 | 0.0111944 | 2.69E-07 |
| $L_8S_8$ | 60 | 1.490 | 0.010 | 0.0111932 | 1.84E-07 |
| $L_8S_8$ | 90 | 1.776 | 0.015 | 0.0111985 | 2.82E-07 |
| $L_8S_8$ | 120 | 1.905 | 0.013 | 0.0112009 | 2.33E-07 |
| $L_8S_8$ | 150 | 1.997 | 0.011 | 0.0112025 | 1.92E-07 |
| $L_8S_8$ | 210 | 1.951 | 0.009 | 0.0112017 | 1.64E-07 |
| $L_8S_8$ | 270 | 1.948 | 0.008 | 0.0112016 | 1.44E-07 |

**Table S1: Results of rubisco KIE assay.** Experimental outputs of rubisco KIE assay; $\delta^{13}C$ vs. time is plotted in Figure S1A. Average $\delta^{13}C$ or $^{13}R$ ($n$=10 analytical replicates) is reported with standard error (standard deviation divided by square root of $n$).

| Strain | Form | Phylum | Specificity | KIE | Notes | Specificity Reference | KIE Reference |
|---|---|---|---|---|---|---|---|
| Prochlorococcus marinus MIT9313 | IA | Cyanobacteria | 59.9 ± 7.0 | 24.0 [22.2, 25.6]a | pH 7.5, 25 mM MgCl2, 25C, expressed from E coli | Shih et al. (2016) | Scott et al. (2007) |
| Synechococcus elongatus PCC6301 | IB | Cyanobacteria | 56.1 ± 1.3 | 22.42 ± 2.37b | pH 8.49, 30 mM MgCl2, 22C, expressed from | This paper | This paper |

| | | | | | | | |
|---|---|---|---|---|---|---|---|
| | | | | | E coli | | |
| Synechococcus elongatus PCC6301 | IB | Cyanobacteria | 50.3 ± 2.0 | 25.18 ± 0.31b | pH 8.38, 30 mM MgCl2, 22C, expressed from E coli | Wang et al. (2023) | Wang et al. (2023) |
| Synechococcus elongatus PCC6301 | IB | Cyanobacteria | 42.7 ± 2.8 | 22.0 ± 0.2c | pH 8.1, 25 mM Mg(2+), 25C, expressed from E coli | Davidi et al. (2020) | Guy et al. 1993 Plant Physiol |
| Candidatus Promineofilum breve | I' | Chloroflexi | 36.1 ± 0.9 | 16.25 ± 1.36b | pH 8.52, 30 mM MgCl2, 22C, expressed from E coli | This paper | This paper |
| Rhodospirillum rubrum | II | Proteobacteria | 12.5 ± 0.6 | 23.0 ± 0.6c | pH 7.9, 25 mM Mg(2+), 25C, expressed from E coli | Davidi et al. (2020) | Guy et al. 1993 Plant Physiol |
| Rhodospirillum rubrum | II | Proteobacteria | 12.5 ± 0.6 | 19.6 ± 0.4c | pH 7.9, 2 mM Mg(2+), 25C, expressed from E coli | Davidi et al. (2020) | Guy et al. 1993 Plant Physiol |
| Rhodospirillum rubrum | II | Proteobacteria | 12.5 ± 0.6 | 22.2 ± 2.1b | pH 8.0, 20 mM MgCl2, room temp?, expressed from E coli (XL1-blue) | Davidi et al. (2020) | McNevin et al (2007) |
| Rhodospirillum rubrum | II | Proteobacteria | 12.5 ± 0.6 | 17.8 ± 0.8b | pH 7.8, 10 mM MgCl2, 25C, "gift from John Schloss" | Davidi et al. (2020) | Roeske & O'Leary (1985) |

**Table S2: Literature compilation of data used to make Figure 1A.** For KIE measurements: Each figure reports uncertainty on the measurement in a different way; superscripts indicate: *a* 95% confidence interval; *b* standard deviation; *c* standard error. Strains for *R. rubrum* not specified in (25, 44, 45). We only used data where a pure enzyme, substrate-depletion assay like ours was done. In addition, we only used data from well-characterized strains where rubisco was obtained through expression in *E. coli*. Therefore, we are not including the (47) measurement because it was done in a tobacco plant mutant expressing an *R. rubrum* rubisco sequence *in vivo*, and KIE was calculated by extrapolating to a ratio of intercellular to ambient $CO_2$ ($C_i/C_a$) of 1. In addition, we do not include the Form II measurement from (48) because the rubisco from a *Riftia pachyptila* endosymbiont was purified from the host trophosome; the endosymbiont is not well characterized because it cannot be cultured separately from the host currently though the complete and closed genome has recently been presented (57). In addition, we are only showing Form IA/B data from Cyanobacteria and therefore do not include plants or the *Solemya velum* symbiont (35). Assay temperature was assumed to be room temperature for (44); rubisco was assumed to be expressed from *E. coli* in (45). In addition, only the pH 7.9, 25 mM $Mg^{2+}$ condition from (25) was plotted in Figure 1B. See Table 3 in (55) for a recent compilation of all measured KIEs. For Specificity measurements: Most specificity values were not reported with the study, with the exception of this paper and (46). Therefore, specificity values were taken from (56, 58) where indicated.

| **Strain** | **Form** | **Specificity** | **KIE (‰)** | **Specificity Reference** | **KIE Reference** |
|---|---|---|---|---|---|
| Ralstonia eutropha | IC | 75 | 19 [17.5, | Horken & Tabita | Thomas et al. |

| | | | | | |
|---|---|---|---|---|---|
| | | | 20.4] | (1999) | (2018) |
| Rhodobacter sphaeroides | IC | 60 | 22.4 [21.1, 24.0] | Horken & Tabita (1999) | Thomas et al. (2018) |
| Emiliania huxleyi | ID | 79 | 11.1 [9.8, 12.6] | Badger et al. (1998) | Boller et al. (2011) |
| Skeletonema costatum | ID | 72.2 ± 2.2 | 18.5 [17.0, 19.9] | Haslam et al. (2005) | Boller et al. (2015) |
| Spinacia oleracea | IB | 77.2 ± 1.4 | 30.3 ± 0.8 | Read and Tabita (1994) | Guy et al. (1993) |
| Spinacia oleracea | IB | 77.2 ± 1.4 | 29 ± 1 | Read and Tabita (1994) | Roeske and O'Leary (1984) |
| Spinacia oleracea | IB | 77.2 ± 1.4 | 28.2 [26.6, 29.8] | Read and Tabita (1994) | Scott et al. (2004) |
| Nicotiana tabacum | IB | 80 | 27.4 ± 0.9 | Kane et al. (1994) | McNevin et al. (2007) |
| Riftia pachyptila symbiont | II | 8.6 ± 0.9 | 19.5 ± 1.0 | Robinson et al. (2003) | Robinson et al. (2003) |

**Table S3: Additional Specificity and KIE values used for Figure 1B.** Data compilation is similar to that used in Figure 4 from (36). Most specificity values were measured separate from the KIE and are taken from other prior literature (59–63), similar to what was done by (36). (48) reports both specificity and KIE. The *Solemya velum* gill symbiont (Form IA, KIE = 24.4‰) from (35) was not included because the specificity could not be found. In addition, (25) gives two values at two different assay conditions for *S. oleracea*; here we use the value at pH 8.5, 20 mM $MgCl_2$ but they also report a KIE of 29.0 ± 0.3‰ at pH 7.6, 5 mM $Mg^{2+}$. KIE values are from (25, 36, 44, 48, 64–66). Error in brackets is reported as mean with 95% confidence intervals; otherwise error is reported as mean ± s.e. We could not get access to (63) so the specificity value for *N. tabacum* (highlighted in orange) is estimated to be ~80 from Figure 4 in (36).

| Rubisco | Parameter | Estimate | Std. Error | t value | Pr(>\|t\|) | Signif. Code |
|---|---|---|---|---|---|---|
| $L_8S_8$ | a | -2.786E-05 | 2.759E-06 | -10.097 | 1.63E-04 | *** |
| $L_8S_8$ | b | 1.640E-02 | 4.310E-03 | 3.804 | 0.012573 | * |
| $L_8S_8$ | c | 1.120E-02 | 2.908E-06 | 3852.671 | < 2e-16 | *** |
| $L_8$ | a | -2.338E-05 | 1.369E-06 | -17.072 | 2.58E-06 | *** |
| $L_8$ | b | 1.769E-02 | 2.767E-03 | 6.392 | 6.90E-04 | *** |
| $L_8$ | c | 1.118E-02 | 1.203E-06 | 9291.937 | < 2e-16 | *** |

**Table S4: Model outputs for converting time to *f*.** The nonlinear least squares function in R Statistical Software was used for calculation with initial guesses of a=-1E-5, b=0.1, c=0.01 for $L_8S_8$; a=-1E-4,b=0.1,c=0.01 for $L_8$. The parameter *c* gives $R_{upper}$ in Equation 1, which then allows time to to be convert to *f*. For $L_8S_8$, the model found a residual standard error of 1.252-06 on 6 degrees of freedom, required 5 iterations to convergence, and achieved a convergence tolerance of 1.405e-06. For $L_8$, the model found a residual standard error of 1.252e-06 on 6 degrees of freedom, required 5 iterations to convergence, and achieved a convergence tolerance of 1.405e-06. The significant codes indicate: 0 ‘***’ 0.001 ‘**’ 0.01 ‘*’

0.05 ‘.’ 0.1 ‘ ’ 1. All analyses were performed using R Statistical Software (v4.1.0; R Core Team 2021, (40)).

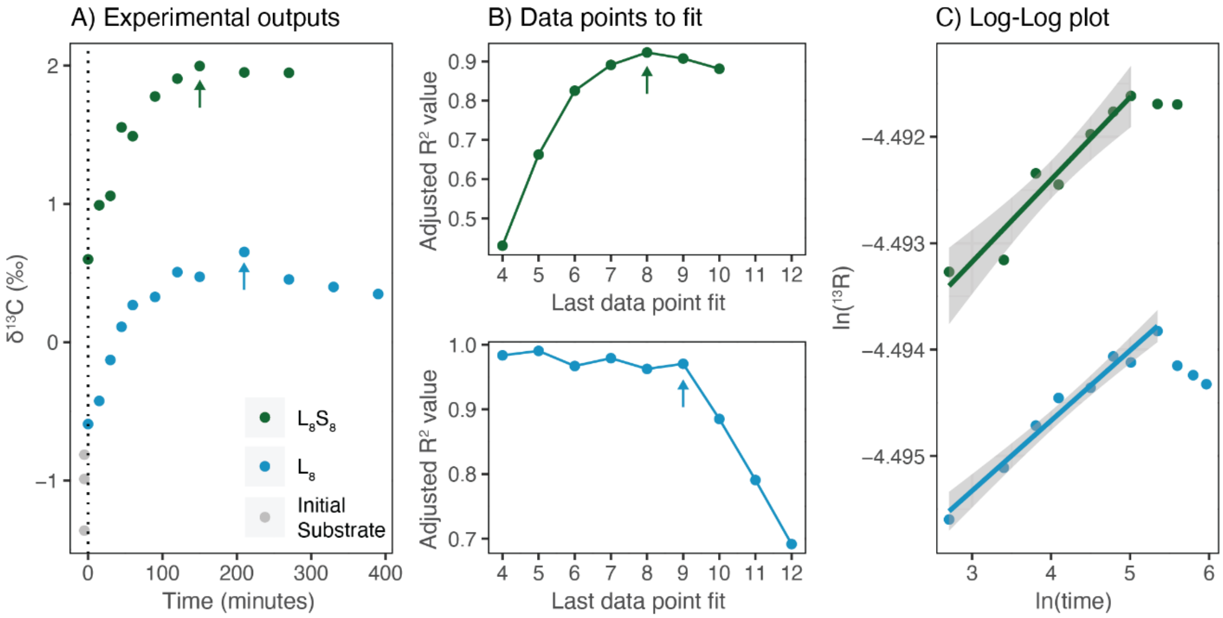


**Figure S1: Data preprocessing step.** A) Experimental outputs of rubisco KIE assay, showing how the $\delta^{13}C$ of the $CO_2$ headspace evolves over the experiment. The first time point taken is shown at 0 minutes, and the initial $NaHCO_3$ substrate is shown plotted at -5 minutes for ease of comparison. The arrow indicates the final data point to fit after preprocessing. B) Subplots showing the adjusted $R^2$ value for the $L_8S_8$ (above, green) and the $L_8$ rubisco (below, blue) for linear regressions across different lengths of log-transformed data points. Arrows indicate where the $R^2$ value starts to decrease (point 8 for $L_8S_8$ rubisco in green; point 9 for $L_8$ rubisco in blue); these arrows refer to the same point in Panel A. C) Linear regression across natural log-transformed data to data point 8 for the $L_8S_8$ rubisco (green) and to data point 9 for the $L_8$ rubisco (blue). Note the isotopic data is in $^{13}R$ vs. $\delta^{13}C$ format. The first data point is not plotted because the natural log of zero is undefined. All analyses were performed using R Statistical Software (v4.1.0; R Core Team 2021, (40)). Data visualization was performed using the ggplot2 package (v3.3.6; Wickham, 2016, (67)).

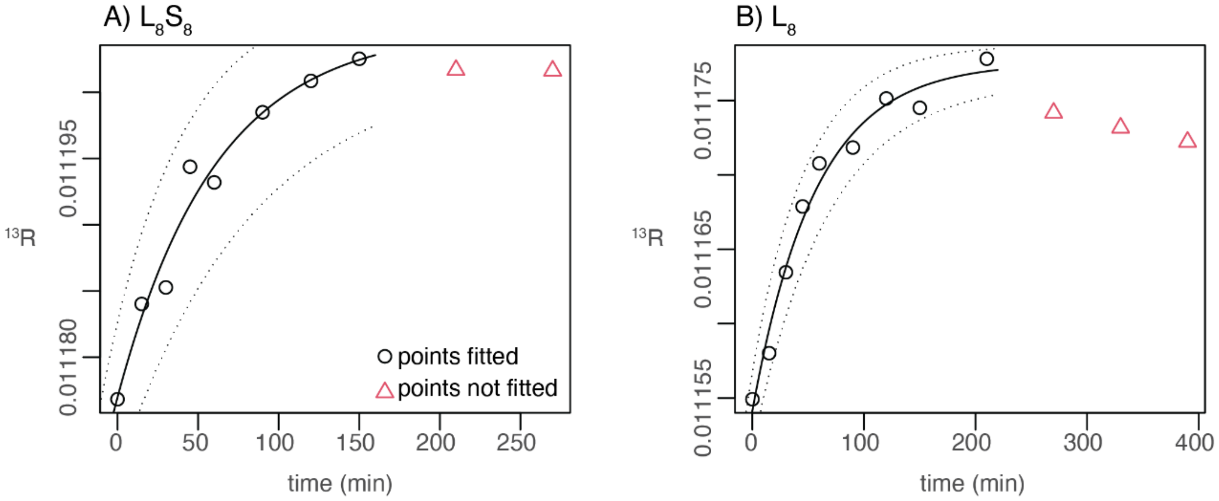

**Figure S2: Calculating *f* from time.** Plots showing best fit exponential model for A) L8S8 vs. B) L8 rubisco in solid black line. Dotted lines indicate model uncertainty (std. dev.). See Table S3 for best-fit model parameters. Open black circles are points fitted, as determined in Figure S1. Open red triangles are the points not fit. All analyses and data visualization were performed using R Statistical Software (v4.1.0; R Core Team 2021, (40)).

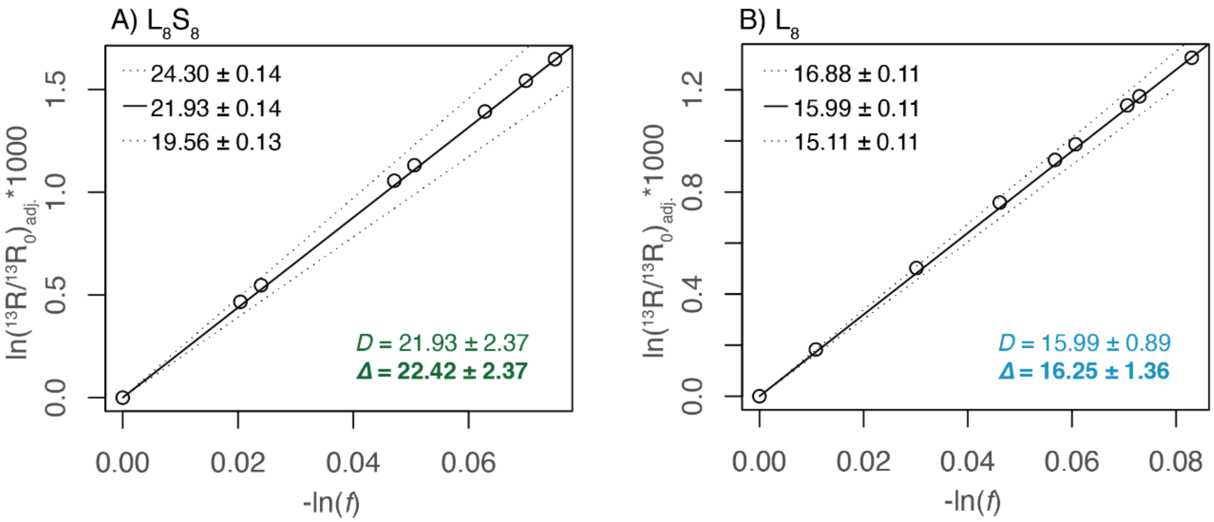


**Figure S3: Rayleigh plots with and without equilibrium adjustment.** A) and B) show $L_8S_8$ and $L_8$ rubisco with equilibrium adjustment for $^{13}R$ values (Equation 2; (25) ) before linear regression. Solid line gives best fit value using *f* values calculated from the best estimate for parameter *c*. Dotted lines give fit for *f* values calculated using the best estimate ± std. error for *c* as shown in Table S3. Slopes for each line are reported in the upper left corner (best estimate ± std. error). *D* is the slope of the solid, best fit line. Δ is converted from *D* using Δ = *D*/(1-*D*/1000) (25). All analyses and data visualization were performed using R Statistical Software (v4.1.0; R Core Team 2021, (40)).


### Funding

R.Z.W. was supported by the National Science Foundation Graduate Research Fellowship Program (NSF GRFP). A.K.L., D.M.B., and P.M.S. were supported by a Society in Science–Branco Weiss fellowship from ETH Zürich and a Packard Fellowship from the David Lucile Packard Foundation. W.F.S. and research was supported by NASA Exobiology (80NSSC21K0484), Simons Foundation Collaboration on Origin and Evolution of Life, Schwartz Reisman Collaborative Science Program, Caltech Center for Evolutionary Sciences.

### Acknowledgements

None


### Author contributions

Conceptualization, W.F.S. and P.M.S; Methodology, R.Z.W., A.K.L., and D.M.B.; Formal Analysis, R.Z.W.; Writing - Original Draft Preparation, R.Z.W.; Writing - Review and Editing, R.Z.W., W.F.S., A.K.L., and P.M.S.; Funding Acquisition, W.F.S. and P.M.S.
